# Dung beetles as vertebrate samplers – a test of high throughput analysis of dung beetle iDNA

**DOI:** 10.1101/2021.02.10.430568

**Authors:** Rosie Drinkwater, Elizabeth L. Clare, Arthur Y. C. Chung, Stephen J. Rossiter, Eleanor M. Slade

## Abstract

The application of environmental DNA (eDNA) sampling in biodiversity surveys has gained widespread acceptance, especially in aquatic systems where free eDNA can be readily collected by filtering water. In terrestrial systems, eDNA-based approaches for assaying vertebrate biodiversity have tended to rely on blood-feeding invertebrates, including leeches and mosquitoes (termed invertebrate-derived DNA or iDNA). However, a key limitation of using blood-feeding taxa as samplers is that they are difficult to trap, and, in the case of leeches, are highly restricted to humid forest ecosystems. Dung beetles (superfamily Scarabaeoidea) feed on the faecal matter of terrestrial vertebrates and offer several potential benefits over blood-feeding invertebrates as samplers of vertebrate DNA. Importantly, these beetles can be easily captured in large numbers using simple, inexpensive baited traps; are globally distributed; and also occur in a wide range of biomes, allowing mammal diversity to be compared across habitats. In this exploratory study, we test the potential utility of dung beetles as vertebrate samplers by sequencing the mammal DNA contained within their guts. First, using a controlled feeding experiment, we show that mammalian DNA can be retrieved from the guts of large dung beetles (*Catharsius renaudpauliani*) for up to 10 hours after feeding. Second, by combining high-throughput sequencing of a multi-species assemblage of dung beetles with PCR replicates, we show that multiple mammal taxa can be identified with high confidence. By providing preliminary evidence that dung beetles can be used as a source of mammal DNA, our study highlights the potential for this widespread group to be used in future biodiversity monitoring surveys.

## Introduction

The development and application of molecular techniques to sequence the DNA contained within environmental samples (eDNA), including water and soil, has provided new opportunities for assaying biodiversity (Beng *et al*. 2016, Robson *et al*. 2016, Nguyen *et al*. 2020). In terrestrial systems, several recent eDNA-based studies aimed at assaying vertebrate biodiversity have sequenced dietary DNA contained within the blood meals of invertebrates (iDNA) (Schnell *et al*. 2018, Fahmy *et al*. 2019), including flies (Gogarten *et al*. 2019), leeches (Tilker *et al*. 2020), and mosquitoes (Kocher *et al*. 2017).

A key limitation of using blood-feeding taxa as samplers is that they are difficult to trap. When sampling for blood-feeding leeches, collection of individuals can be opportunistic (Schnell *et al*. 2012) or, in more recent studies, by hand-searching within fixed areas (Abrams *et al*. 2019, Drinkwater *et al*. 2020). Additionally, some blood-feeding invertebrate taxa being used in these types of studies, specifically terrestrial leeches, are highly restricted to humid forest ecosystems (Borda & Siddall 2010). For example, the distribution of haemadipsid leeches with the potential for use in iDNA studies extends only across Southeast Asia, India and Madagascar (Schnell *et al*. 2018) and within these regions their occurrence is linked to humid habitats (Drinkwater *et al*. 2019).

Dung beetles (superfamily Scarabaeoidea) are a diverse and wide-ranging group, that feed primarily on the faecal matter of terrestrial vertebrates. As detritivores, dung beetles provide many crucial ecosystem functions and services, such as seed dispersal, nutrient cycling, and greenhouse gas reduction in both tropical and temperate ecosystems (Nichols *et al*. 2008). Two recent studies have demonstrated that the epithelial cells of mammals retained in the dung ingested by these beetles may, like blood meals, provide a viable source of vertebrate DNA. Gómez & Kolokotronis (2016) used Sanger sequencing to recover mammal DNA from the guts of individual dung beetles feeding on horse manure, while Kerley *et al*. (2018) successfully applied metabarcoding to retrieve mammal DNA from individual dung beetle faeces. These early findings were based on the sequencing of individual samples and imply that coprophagous insects may represent promising alternatives to blood-feeding models in iDNA studies. Adult dung beetles gain nutrition from liquid in dung by concentrating microorganisms and vertebrate cells through particle feeding (Nichols & Gómez 2014); therefore, if iDNA is to be detected in the gut, mammal epithelial cells first need to first pass through the dung beetle’s filtering mouth parts. The size of particle that a beetle can ingested has been shown to be size- and species-dependent (Holter *et al*. 2002, Holter & Scholtz 2007), and, as such, we might expect an effect of size on the recovery of iDNA.

As potential samplers, dung beetles offer several distinct advantages over existing blood-feeders. First, they are found in most terrestrial habitats, ranging from temperate zones to the equatorial tropics, and occur on all continents except Antarctica (Nichols & Gardner 2011). Second, dung beetles can be easily captured in large numbers using low-cost home-made traps, allowing for standardised sampling regimes (Nichols & Gardner 2011). In the Brazilian Amazon, dung beetles were identified as one of the most cost-effective taxa for biological surveys and had the highest ecological indicator value (Gardner *et al*. 2008). Third, some dung beetles specialise on the dung of different mammal species or guilds (e.g., Raine & Slade 2019); thus it may be possible to use some species to detect specific mammals of interest. Conversely, by sampling multispecies assemblages of dung beetles, it should be possible to detect a range of mammal species present in an area.

Here we aim to gain a better understanding of the utility of dung beetles as iDNA samplers for biodiversity studies in the humid tropics. To this end, we (i) ascertain the time window over which mammal DNA can persist in the gut and still be recovered for amplification and sequencing. For this, we performed a field-based feeding experiment under controlled conditions, focusing on a large-bodied species, *Catharsius renaudpauliani* (Ochi & Kon), which occurs across Borneo. The duration of time over which mammal DNA can be retrieved after a feeding event has previously been characterised for leeches (Schnell *et al*. 2012) and is likely to be a key parameter in interpreting the results of any iDNA-based biodiversity assessments using dung beetles. Additionally, (ii) we apply a high-throughput DNA extraction and sequencing pipeline to pooled samples of dung beetles of different species and assess whether these multi-species assemblages can be used to assay mammal diversity. As with other iDNA studies, pooling samples before sequencing can potentially increase cost-effectiveness and maximise detection rates.

## Methods

### Field-based feeding experiment

#### Sample collection and gut dissection

To measure the window of detection of mammal DNA within dung beetle guts, we constructed a controlled feeding experiment using individuals of the largest dung beetle species commonly occurring in our study area, *Catharsius renaudpauliani*. We collected 60 *C. renaudpauliani* individuals from standard human dung baited live pitfall traps, which were deployed for 24 hours at multiple locations as part of another study (see Parrett *et al*. 2019 for collection details). These traps were set across a habitat gradient, from selectively logged forest to oil palm plantations within the SAFE landscape, Sabah, Malaysia (see Ewers *et al*. 2011 for a full site description). Individuals were collected and maintained in sex-specific “holding” boxes, with moist sand and *ad libitum* cow dung for three days. Cow dung was used as the only non-human dung that could be obtained in bulk from nearby farms.

To measure the persistence of cow DNA in beetle guts, individual beetles were transferred to clean enclosures and starved for 48 hours to purge any dung from their guts. Twenty grams of cow dung was then introduced into the boxes and the beetles were left to feed *ad libitum* for approximately one hour, to allow all individuals the opportunity to feed. Remaining dung was then removed, and the enclosures were cleaned thoroughly. At 10 set time points (0, 1, 2, 4, 6, 9, 12, 24, 48, 56 hours post feeding) six individual beetles (three females and three males) were selected *ad hoc* and frozen for at least an hour before decapitation. The length of the beetle (a proxy for beetle size) was recorded using callipers before the guts were dissected under sterile conditions placed in 3-4 times the gut volume of RNALater and stored in a freezer for DNA preservation. Very little is known of metabolism and digestion in this species, but rapid digestion of the dung was assumed, following Upadhyay (1983) who found that full digestion occurred within 48 hours in a congeneric species. We therefore used 48 hours as the purging time and we aimed to maximise the number of early time points sampled post-feeding to capture patterns of DNA degradation.

#### Quantification of DNA

DNA was extracted from all beetle guts applying the same protocol used in Drinkwater *et al*. (2018) for the extraction of iDNA from terrestrial leeches. This involved digesting each sample overnight with proteinase K and lysis buffer, then extracting the DNA using a QiaQuick purification kit (Qiagen) following manufacturers protocols with reduced centrifuge speeds (full details in Drinkwater *et al*. 2018). Quantitative PCR (qPCR) was used to determine the concentration of cow DNA detected from the beetle guts. The six DNA extracts, from three female and three male guts, from each time point were initially diluted by a factor of five to improve qPCR efficiency. qPCR reactions were then set up in a total volume of 20μl using SYBR green fluorescence as the marker. Each reaction consisted of 10μl of SensiFAST mastermix (Bioline, UK), 0.8μl of 10μM primers, both forward and reverse, 7.4μl of ddH20, with 1μl of unknown DNA template. 16s rRNA primers were used to target small fragments of mammal DNA (Taylor 1996). For quantification, we used a standard curve of eight samples of known DNA concentrations which were included in the qPCR plate alongside the unknown gut samples (standard curve in figure S1). For confirmation of the identity of the qPCR product a subset of the qPCR reactions were sequenced using Sanger sequencing, and the species identified from the NCBI GenBank database with BLAST. The copy number of the samples was calculated using the equation: 10^(CT – intercept)/slope^, where the intercept and slope are calculated from the standard curve. The effect of time post-feeding on the number of DNA copies recovered was tested using a log-log linear regression model, with log DNA copy number as the response variable, and log time in hours (post-feeding) as the main effect. Additionally, sex and size of the beetle were included in the models.

### Sequencing of iDNA from multi-species assemblages

#### Sample collection

We used two types of human-baited pitfall traps, deployed opportunistically in an area of continuous logged forests for 24 hours. The traps were either traditional pitfall traps, with a ball of dung held in muslin cloth suspended over the trap, used for surveying dung beetle composition (see Slade *et al*. 2011 for methods), or traps where beetles were excluded from the dung ball, using a plastic cup, a precautionary attempt at reducing contamination from human DNA.

From the pitfall traps we sampled either individuals of *Catharsius* spp. or the whole trap assemblage. We dissected the guts of 18 large *Catharsius* spp. beetles in the field. To reduce contamination risk, we performed the dissections under a covered box, ensuring dead air, wiping down surfaces with ethanol, changing gloves, and flaming scalpels. This was an initial exploration as to whether sterile dissections were possible in limited conditions, making iDNA studies more accessible and logistically easier for field ecologists. Each gut was then placed into an individual tube, containing 3-4x the gut volume of RNALater and stored in the field freezer (with approximately 10 hours of power per day). To compare these filed dissections with dissections done under sterile laboratory conditions, we took a further six *Catharsius* spp. individuals from the traps and stored them whole in ethanol for gut dissection in laboratories at Queen Mary University of London, UK. Finally, the entire contents of two traps were stored in ethanol to be sequenced as an assemblage, without prior gut dissection. The 24 individual gut DNA extractions were pooled into 3 pools for sequencing (field dissected guts 2 x 6 individuals, laboratory dissection 1 x 6 individuals). The assemblage traps were split into 3 DNA extraction pools per trap. Resulting in a total of nine pools for amplification and sequencing (Table 1). All DNA extractions were conducted as above (see qPCR DNA extractions) following the extraction and sample pooling protocol used in Drinkwater et al., (2018) at Queen Mary University of London.

**Table 1.**
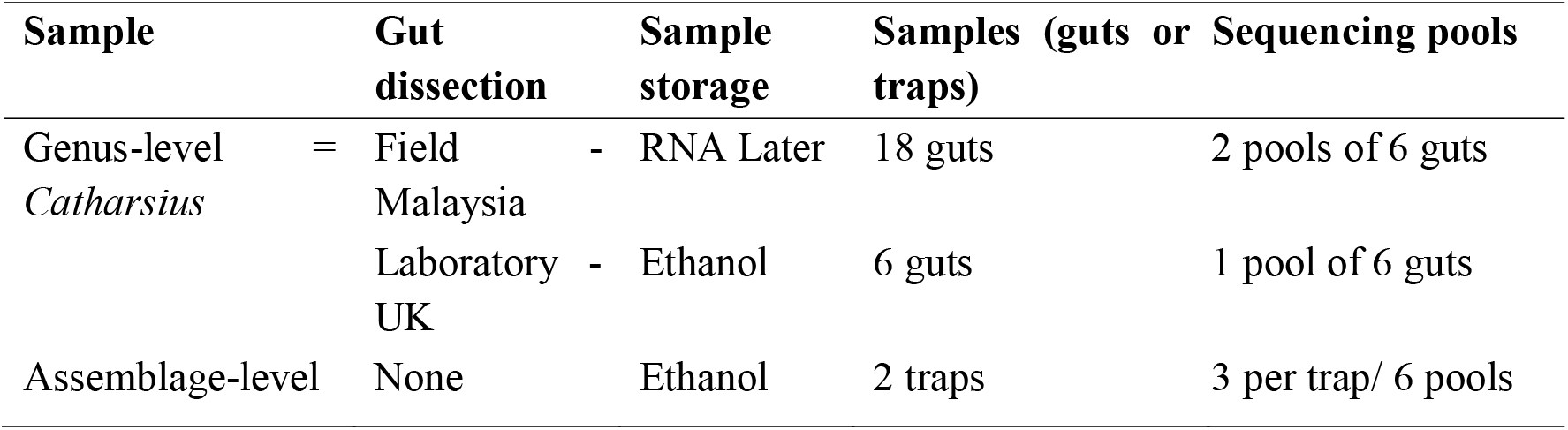
Summary of samples and pooling used in the study

#### PCR amplification and sequencing

We used primers targeting mammalian 16S rRNA, which previous studies have shown to be successful for identifying mammals in leech iDNA (Taylor 1996). Following the laboratory protocols for high throughput sequencing of leech iDNA in Drinkwater *et al*. (2018) each DNA extract was amplified using uniquely tagged primers (Binladen *et al*. 2007) and, extraction blanks and negative PCR controls were included in each PCR run. The reactions consisted of 1μl of template DNA in 0.2mM of 10×buffer, 2.5mM MgCl2, 1 unit DNA polymerase (AmpliTaq Gold, Applied Biosystems), 0.2mM dNTP mix (Invitrogen), 0.5mg/ml BSA, and 0.6μM of the forward and reverse primer to make a final reaction volume of 2μl. We used thermocycling conditions of 95°C for 5min, then 40 cycles of 95°C for 12s, 59°C for 30s and 70°C for 20s with a final extension time of 7min at 70°C. Amplification was checked on a 1% agarose gel, successful reactions were pooled for DNA amplicon libraries (Carøe *et al*. 2017) and subjected to paired end sequencing with 150 bp Illumina MiSeq at The Genome Centre at Queen Mary University of London.

#### Bioinformatics and taxonomic identification

We merged forward and reverse reads with AdapterRemoval version 2 (Schubert *et al*. 2016) and sorted samples by their unique 16s primer tags allowing the identification of the original sample before filtering using DAMe (Zepeda Mendoza *et al*. 2016; following version updates at: https://github.com/shyamsg/DAMe). We filtered based on length using a minimum length cut-off of 90 bp and unpaired reads were removed. We clustered the reads into operational taxonomic units (OTUs) at 97% similarity using SUMACLUST (Mercier *et al*. 2013). OTUs were then checked for chimeras using mothur (Schloss *et al*. 2009) and further filtering of OTUs was conducted using LULU (Frøslev *et al*. 2017). OTUs were identified using a BLAST search against a customised reference database, resulting in a list of taxa for each dung beetle gut iDNA sample. The reference database contained all available 16S mammal sequences for Bornean mammals and known lab contaminants (Table S1). Where reference sequences did not exist for a species, a closely related taxon was included in the database. Due to our small sample size and the exploratory nature of the study, we present the results as descriptive data. Although read count is not a representative measure of detection due to the uneven digestion and amplification (e.g. PCR) processes (Deagle *et al*. 2018), we have included this in the summary table of detections to highlight the potential of DNA recovery (Table 3).

## Results

### Window of DNA persistence in C. renaudpauliani guts

The standard curve shows that the efficiency of the qPCR reactions (R^2^) was greater than 0.99. The results of the log-linear qPCR analysis indicate a decrease in DNA copy number over time (Figure 1). This is supported by results of the simplified log-log linear model containing only (log) time post-feeding (Table 2). The model R^2^ value was 0.65, indicating that 65% of the variance in the DNA copy number could be explained by time. Parameter estimates from this model indicate that DNA copy number decreases with time post feeding (Figure 1, Table 2). We did not find a relationship between the weight (t = −1.842, p 0.07, df = 4/41), sex (t = 0.01, p = 0.99, df = 4/41) or length (t = 0.84, p = 0.40, df = 4/41) of the beetle and these terms were therefore removed.

**Figure 1.**
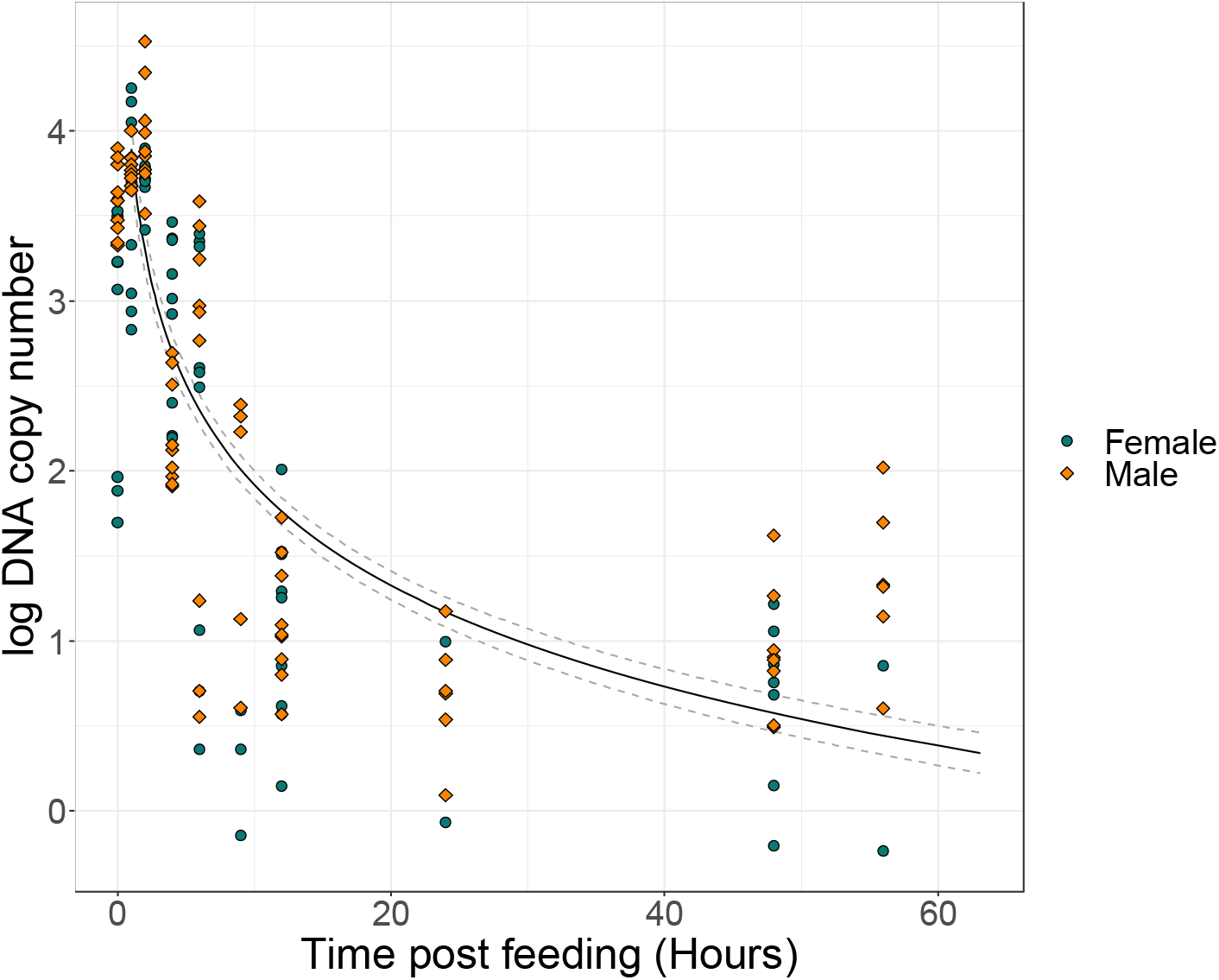
Comparison of log DNA copy number as a function of time post-experimental feeding. The blue circles refer to the samples from female beetles and the orange diamonds to samples from male beetles. The line represents the model fit of the log-log linear model.

**Table 2.** Final model output for the log-log linear model of DNA copy number with time post feeding.

| Coefficient | Estimate | Standard error | t value | p value |
| --- | --- | --- | --- | --- |
| Intercept | 3.79 | 0.12 | 31.07 | <0.05 |
| Log10(Time) | -1.97 | 0.12 | -16.23 | <0.05 |

### Identity of mammal species from dung beetle assemblages

We identified six mammalian taxa in the iDNA from the beetles caught in multi-species assemblage traps (Table 3). These mammals were from five families and represented the common species in the area. The detection rate was just under 50% with four out of the nine pools sequenced resulting in detections. All traps, regardless of the modifications, generated a large amount of human DNA contamination.

**Table 3.**
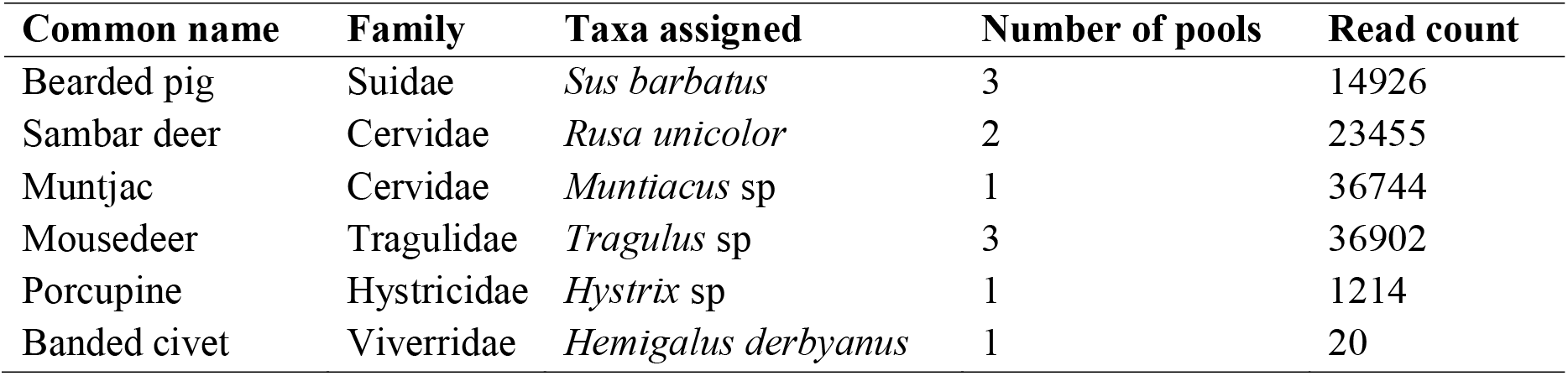
Taxa detected in the dung beetle gut iDNA (either mammal species or genus), the number of pools it was recorded in, and the DNA read count.

## Discussion

In this paper we have demonstrated that iDNA from mammalian sources can be recovered from the guts of tropical forest dung beetles. We achieved this using a high throughput sequencing pipeline, developed for leech-based biodiversity surveys (Drinkwater *et al*. 2018). We found that there was rapid digestion and fast passage of cow dung through beetle guts. The raw values show very high initial DNA copy number up to 2-4 hours followed by a sharp decrease to zero DNA recovery at 9 hours post feeding (Figure S2) and our model showed that there is an approximately 2% decrease in DNA copy for every 1% increase in time.

There has been very little previous work on the digestion of dung in dung beetles, but broadly our finding corroborates that of Upadhyay (1983) who performed feeding observations and also reported a short digestion window of 48 hours in *Catharsius molossus*, a member of the same genus of large dung beetle. This is in contrast to the blood feeding leeches *(Hirudo medicinalis)*, for which Schnell *et al*. (2012) found that iDNA could be detected for up to four months. The marked difference in the time window of detection offered by dung beetles, highlights the potential benefit of combining these two invertebrate samplers to target mammal diversity. At the same time, however, ours is a preliminary experiment conducted under field conditions in Borneo, in which cow dung was used for both the pre- and post-feeding. For this reason, we cannot rule out the possibility that cow DNA detected post-feeding could have persisted from a previous feeding event, although we experimental procedure was designed to avoid this. Indeed, we did not detect any DNA ~20 hours post-feeding and the beetles were given a 48-hour purging window once they had been exposed to the cow dung; thus, we believe that the detected DNA was the target DNA from our experimental feeding.

Our results revealed no relationship between DNA yield and gut weight, which was supervising given that we would expect heavier guts to contain more contents and, therefore, more iDNA. Additionally, we did not find a relationship between DNA copy number and beetle length (a proxy for size). As adult dung beetles are filter feeders, we would expect the detection rate to be associated with size, as to sequence iDNA the epithelial cells from the dung source need to be able to pass through the beetle epipharynx (Holter & Scholtz 2007). Intra-specific variation within the *Catharsius* individuals we sampled may not have been variable enough to demonstrate any impact of size. However, as this is the known mechanism for feeding in adult beetles, it could be beneficial to repeat the experiment using species with a wider range of variation in body size.

Our assays of multi-species beetle assemblages led to detection of six mammalian taxa, representing five families. Three of these could be resolved to species level, whereas three could only be confidently identified to genus, as there are two congeneric species present across the site. Additionally, given that we have now demonstrated a possible temporal threshold of DNA persistence in guts of *C. renaudpauliani*, the results suggest that when mammal DNA is detected that feeding is most likely to have occurred within four hours of being trapped. Dung beetles are attracted to fresh dung, which is removed quickly in tropical forests (with even large dung piles completely removed within 24 hours) (Slade *et al*. 2011).

Our findings therefore suggest that the mammals detected by iDNA were occupying the area within the temporal window of the trapping campaign. Although this requires further research, the potential to “time-stamp” iDNA detections in this way could be beneficial for conservation applications.

The most frequently detected mammals were the most common and larger bodied species. This indicates that ungulate species found in the region, such as the bearded pig, muntjac and sambar deer, may be a key dietary resource for dung beetles. In addition, *Hystrix sp* was also positively identified and could be assigned to one of two *Hystrix* species on Borneo, the endemic thick-spined porcupine or the Malay porcupine, both of which are relatively large and abundant. We also recorded the banded civet from one trap, which is a species of conservation concern due to declining population trends and is listed as near threatened on the IUCN red list (Ross *et al*. 2015). The presence of all these species have been confirmed in the area using leech-based iDNA sampling (Drinkwater *et al*. 2020). Taking these results together, our proof-of concept study clearly highlights the usefulness of combining multiple iDNA samplers, which offer the potential of targeting two different windows of detection, one short term (i.e., beetles) and one longer term (i.e., leeches). We also note that a high amount of human DNA was recovered even when using the most sterile techniques. Although some of this DNA will have arisen through laboratory or field contamination, it is likely that it may also represent true feeding events. In particular, our study was conducted in a modified landscape consisting of logged forest and oil palm agriculture, with associated human settlements and industrial infrastructure alongside a research field station. Humans could therefore represent an abundant and consistent food source for the dung beetles in this area.

Previous studies in South Africa have detected DNA from common and cryptic mammal species using shot-gun sequencing of multi-species assemblages (Gillett *et al*. 2016) and metabarcoding of iDNA from a single dung beetle species Kerley *et al*. (2018). The speed and cost-effectiveness of the field sampling using dung beetles, means that it could be beneficial to use dung beetle iDNA surveys alongside comprehensive camera trapping surveys to supplement detection data. The validation of iDNA surveys compared to camera trapping is an active area of research. The low field input of leech iDNA compared to camera trapping has been highlight before (Weiskopf *et al*. 2017). Now the focus is moving towards the development of standardised invertebrate collection methods and biodiversity analyses (Abrams *et al*. 2019, Drinkwater *et al*. 2020) allowing for greater integration of the two techniques (Tilker *et al*. 2020). Studies have shown that by combining the results of iDNA with camera traps, and using an occupancy modelling framework can increase the confidence in the estimates, therefore making the results more relevant to wild-life monitoring programmes (Abrams *et al*. 2019). In Laos and Vietnam, a combination of camera trapping and leech iDNA has been used to produce spatial maps for identifying priority areas for conservation (Tilker *et al*. 2020).

Although we mainly focused on *Catharsius*, as the largest beetles in the area, different species of dung beetles have been shown to feed on different mammal dung types (Raine & Slade 2019), and so using mixed species assemblages is likely to be the best approach if using dung beetle samplers to assess the diversity of mammals in an area. *Catharsius*, however, are primarily nocturnal, and as such may feed primarily on the larger mammal dung of nocturnal animals, such as pigs, which could explain the patterns we find in the detections. As well as not recovering detections from small mammals, we also did not detect primates which again may be partially explained by the sampler species choice. The smaller, diurnal beetles in the genus *Onthophagus*, are thought to feed more on the diurnal primate dung (Slade E., pers. comm). We also found that the only detection of a banded civet was in the community trap sample, which consisted of the smaller dung beetles. This may indicate a difference in the diets of the smaller beetles, however, we would need further studies which utilise multiple beetle species, to test robustly whether they capture a different subset of the vertebrate community.

While further work is needed to assess the utility of dung beetles as iDNA samplers, our preliminary data suggest that they may have clear benefits over other invertebrate samplers for conducting low-cost standardised surveys across large areas. Notably, dung beetles occur across a wide range of biomes, and the potentially short gut retention time means the source location of any detected mammal can be more easily placed. Dung beetles are also a bioindicator taxon (Gardner *et al*. 2008), meaning there is the potential opportunity to use iDNA as a way to build quantitative networks of interactions between individual dung beetle and mammal species. Such networks have been attempted using traps baited with different mammal dung types (Raine *et al*. 2019, Ong *et al*. 2020), but these networks only show indirect interactions through the of attractiveness of dung to the beetles, rather than showing direct feeding interactions. By elucidating these direct interactions targeted dung beetle community surveys could be used to assess the health of mammal communities.

## Acknowledgements

This study was funded by NERC Human Modified Tropical Forests program, and we would like to thank the following organisations: Sabah Biodiversity Council, Yayasan Sabah, the Sabah Forestry Department, and Benta Wawasan, for access to field sites, the Stability of Altered Forest Ecosystems (SAFE) project, Sime Darby, and the South East Asian Rainforest Research Partnership (SEARRP) for support in the field. We thank Prof Dr Henry Bernard for logistics and assistance with access, Joseph Williamson, Jonathan Parrett, Sabidee Mohd.Rizan and Herry Heroin, for setting traps and Thomas Gilbert, Ida Schnell and Kristine Bohmann for guidance and assistance in the laboratory.

## Permits

Access and export permits to RD and EMS - JKM/MBS.1000□2/2 (34) JKM/MBS.1000□2/3 JLD.2 (107) and JKM/MBS.1000□2/3 JLD.3 (44))

## Data accessibility statement

Data is available on the SAFE project Zenodo repository XXXXXXXXX. Raw sequence data is available on NCBI short read archive (bioproject accession number pending).

