## Supplemental figure 1 for "Dung beetles as vertebrate samplers – a test of high throughput analysis of dung beetle iDNA"

**Supplementary figures**

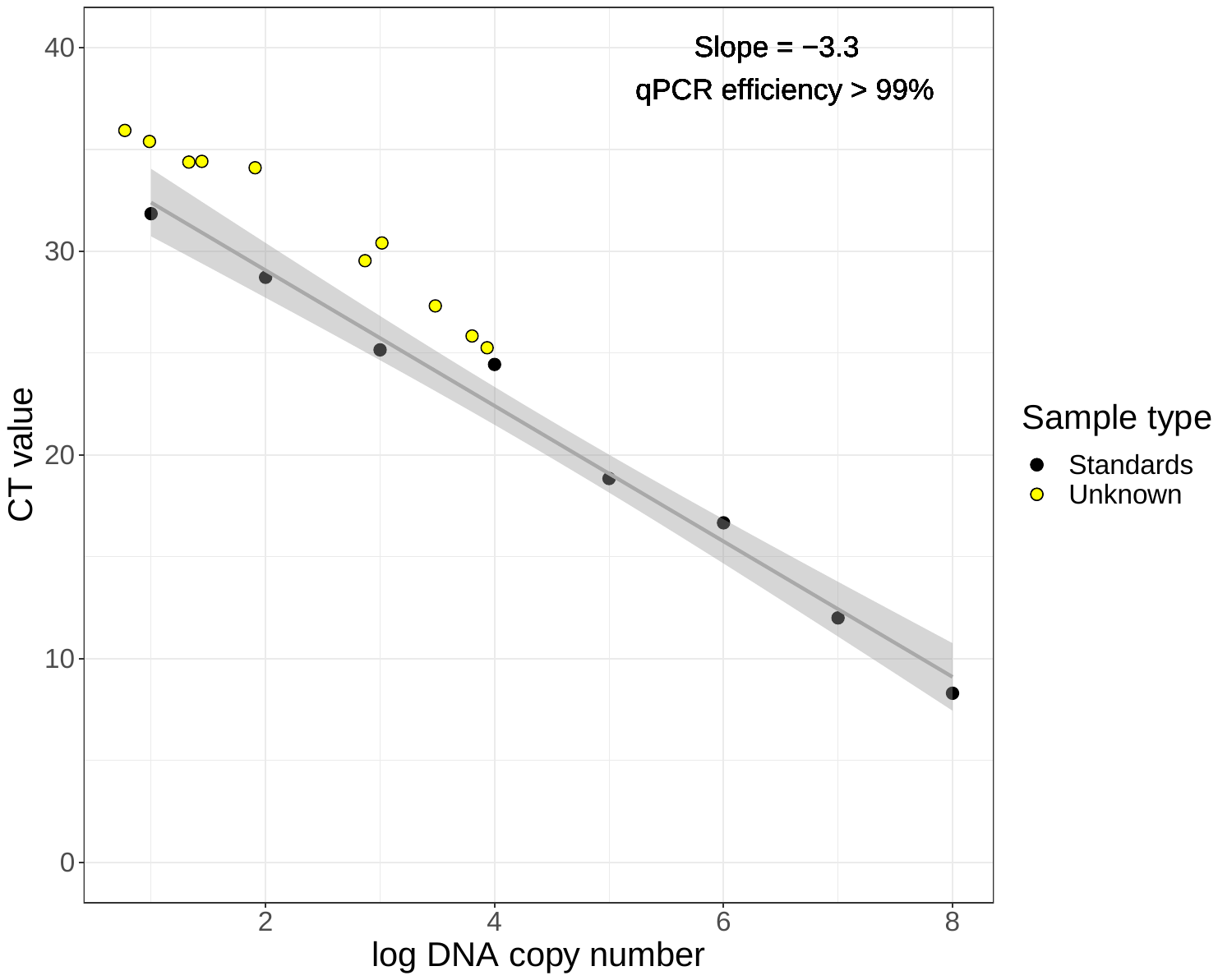

**Figure S1 – qPCR standard curve with an efficiency of 99%.** In black are the CT values of each standard of known concentration. Using the slope of the line (where slope = -3.3 and intercept = 35.72) the efficiency of the reaction is calculated as 99.76% using the standard equation E = -1+10^(-1/slope)^. For qPCR, the desired range of efficiency is between 90-110%. Yellow points refer to the mean CT values for the gut samples at each time point post feeding.
