## Supplemental figure 2 for "Dung beetles as vertebrate samplers – a test of high throughput analysis of dung beetle iDNA"

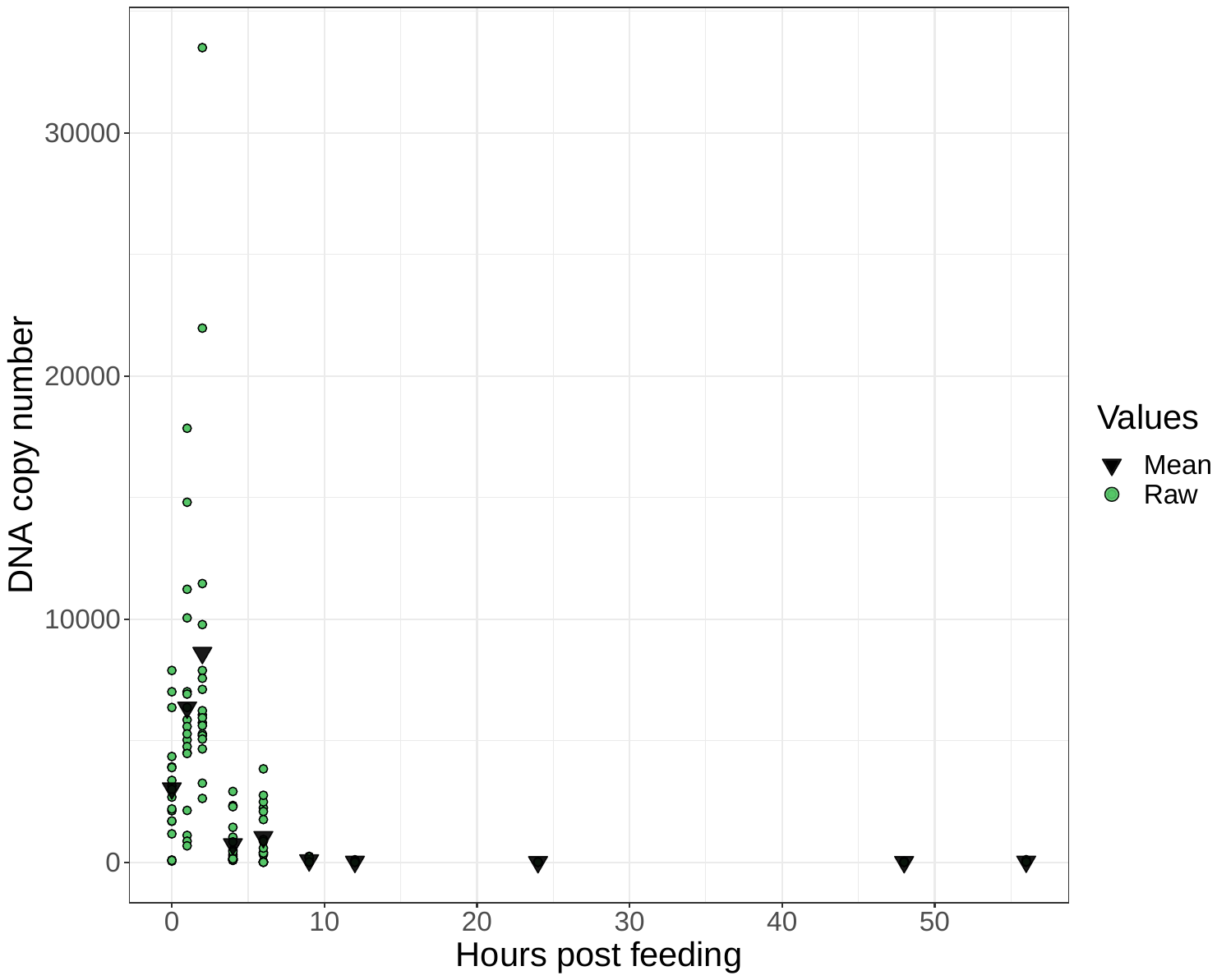

**Figure S2** – Raw values of DNA copy number for each sample at each time point post-feeding. Green points show the raw DNA copy number and black diamonds are the mean value for each time point.
